# Testing for functional organization of three-dimensional surface tilt encoding within visual cortex

**DOI:** 10.1101/2020.01.15.907683

**Authors:** Reuben Rideaux, Andrew E Welchman

## Abstract

Visual perception of three-dimensional (3D) structure is important for object recognition, grasping, and manipulation. The 3D structure of a surface can be defined in terms of its slant and tilt. Previous work has shown that slant and tilt are represented in the posterior and ventral intraparietal sulcus of the human brain; however, it is unclear whether the representation of these features is functionally organized within this region. Here we use phase-encoded presentation of 3D planar surfaces with linear gradients defined by horizontal binocular disparity while measuring fMRI activity to test whether the representation of 3D surface tilt is functionally organized within visual cortex. We find functionally defined structures within V3A and V7. Most notably, in one participant we find that the tilt preference is unilaterally organized in a pinwheel-like structure, similar to those observed for orientation preference in V1, which encompasses most of area V3A. These findings indicate that 3D orientation is functionally organized within the human visual cortex, and the evidence suggesting the presence of a large pinwheel-like structure indicates that this type of organization may be applied canonically within the brain at multiple scales.

## INTRODUCTION

Visual perception of three-dimensional (3D) structure is important for object recognition, grasping, and manipulation. Binocular disparity (i.e., the horizontal offset between the images seen by the left and right eyes) is a powerful cue from which 3D structure can be inferred. Considerable imaging work has progressed our understanding of the basic processing of binocular disparity information is processed in the human brain^1–6^; however, less is known about the regions that encode 3D orientation (parameterized here as slant and tilt).

Electrophysiological work has identified areas in monkey cortex that contain populations of neurons tuned to 3D orientation defined by binocular disparity, including the caudal bank of intraparietal sulcus^7–9^ (CIP) and middle temporal area^10^ (MT). These neurons encode 3D orientation; however, whereas the response of neurons in CIP to slant and tilt is co-dependent^9^, these dimensions appear to be encoded separably in MT^10^. This work has revealed how 3D orientation is encoded at the single-cell level, but the nature of the technique precludes identification of population level trends such as functional organization within these areas. Further, differences between monkey and human brain organization in relation to the representation of 3D orientation restricts the application of these findings to human perception^11^. Here we define functional organization as any non-arbitrary representation of 3D orientation within one or more localized cortical regions.

Functional MRI work with humans has shown that the posterior and ventral intraparietal sulcus (PIPS/V3A & VIPS/V7, respectively) encode depth structure from binocular disparity^12–14^. These areas have also been implicated in processing 3D orientation defined by motion^11,15,16^, texture^17^, and shading^18^. Thus far, the study of how 3D orientation is represented within the human brain has been limited to identification of regions through contrasting patterns of fMRI activation. This technique has been successfully applied in the identification of brain areas that encode 3D orientation, but it has not revealed whether neural responses in these areas are functionally organized.

Here we use phase-encoded presentation of 3D stimuli while recording repeated in-depth measurements of fMRI activity from four human participants to test whether the representation of 3D orientation (i.e., tilt) is functionally organized within visual cortex. We find functionally defined structures within V3A and V7. Most notably, in one participant we find that tilt preference is unilaterally organized in a pinwheel-like structure, similar to those observed for orientation preference in V1^19,20^, that encompasses most of area V3A. Further, for two participants we find an abrupt reversal in tilt preference at the (retinotopically defined) border between V3A and V7. These findings show that it is possible to use non-invasive imaging to discover the functional organisation of 3D orientation within the human visual cortex. Observing pinwheel-like structures suggests that this type of organization may be applied canonically within the brain at multiple scales.

## METHODS

### Participants

Four healthy participants from the University of Cambridge with normal or corrected-to-normal vision participated in the experiment. With the exception of participant 1 (author RR), participants were naïve to the aims of the experiment. The participants ages were 30, 28, 21, and 29, respectively; all were right-handed and participant 3 was female. Participants were screened for stereoacuity using a discrimination task in which they judged the (near/far) depth profile of a random-dot stereogram (RDS) depicting an annulus surrounding a disk. The difference in depth between the annulus and disk was controlled using a 2Down 1-up staircase procedure and participants were admitted into the experiment if they achieved a threshold of <1° arcmin. Participants were also screened for contraindications to MRI prior to the experiment. All experiments were conducted in accordance with the ethical guidelines of the Declaration of Helsinki and were approved by the University of Cambridge STEM and all participants provided informed consent.

### Apparatus and stimuli

Stimuli were programmed and presented in MATLAB (The MathWorks, Natick, MA) with Psychophysics Toolbox extensions^21,22^. Stereoscopic presentation in the scanner was achieved using a “PROPixx” DLP LED projector (VPixx Technologies) with a refresh rate of 120 Hz and resolution of 1920 × 1080, operating in RB3D mode. The left and right images were separated by a fast-switching circular polarization modulator in front of the projector lens (DepthQ; Lightspeed Design). The onset of each orthogonal polarization was synchronized with the video refresh, enabling interleaved rates of 60 Hz for each eye’s image. MR-safe circular polarization filter glasses were worn by participants in the scanner to dissociate the left and right eye’s view of the image. Stimuli were back-projected onto a polarization-preserving screen (Stewart Filmscreen, model 150) inside the bore of the magnet and viewed via a front-surfaced mirror attached to the head coil and angled at 45° above the participants’ heads. This resulted in a viewing distance of 72 cm, from which all stimuli were visible within the binocular field of view. Stereoscopic presentation out of the scanner was achieved using a pair of Samsung 2233RZ LCD monitors (120 Hz, 1680×1050) viewed through mirrors in a Wheatstone stereoscope configuration. The viewing distance was 50 cm and participants’ head position was stabilized using an eye mask, head rest and chin rest. Eye movement was recorded binocularly at 1 kHz using an EyeLink 1000 (SR Research Ltd., Ontario, Canada).

Stimuli consisted of RDS presented within an annulus (inner radius, .45°; outer radius, 9°) aperture on a mid-grey background surrounded by pink noise intended to facilitate stable vergence. The outer edge of the annulus was blurred according to a cosine profile. Dots in the stereogram followed a black or white Gaussian luminance profile, subtending 0.17° at half maximum. There were 12 dots/deg_2_, resulting in ~85% coverage of the background. Dots were allowed to overlap and where this occurred, they occluded previously positioned dots. Dots presented to the left and right eyes were offset to generate planar surfaces with linear gradients of horizontal disparity (**Fig. 1a**). Binocular disparity was calculated from the cyclopean view and applied to each dot based on the specific disparity-defined slant/tilt angle. In the centre of the annulus, we presented a fixation square (side length = 0.5°) paired with horizontal and vertical lines.

**Figure 1.**
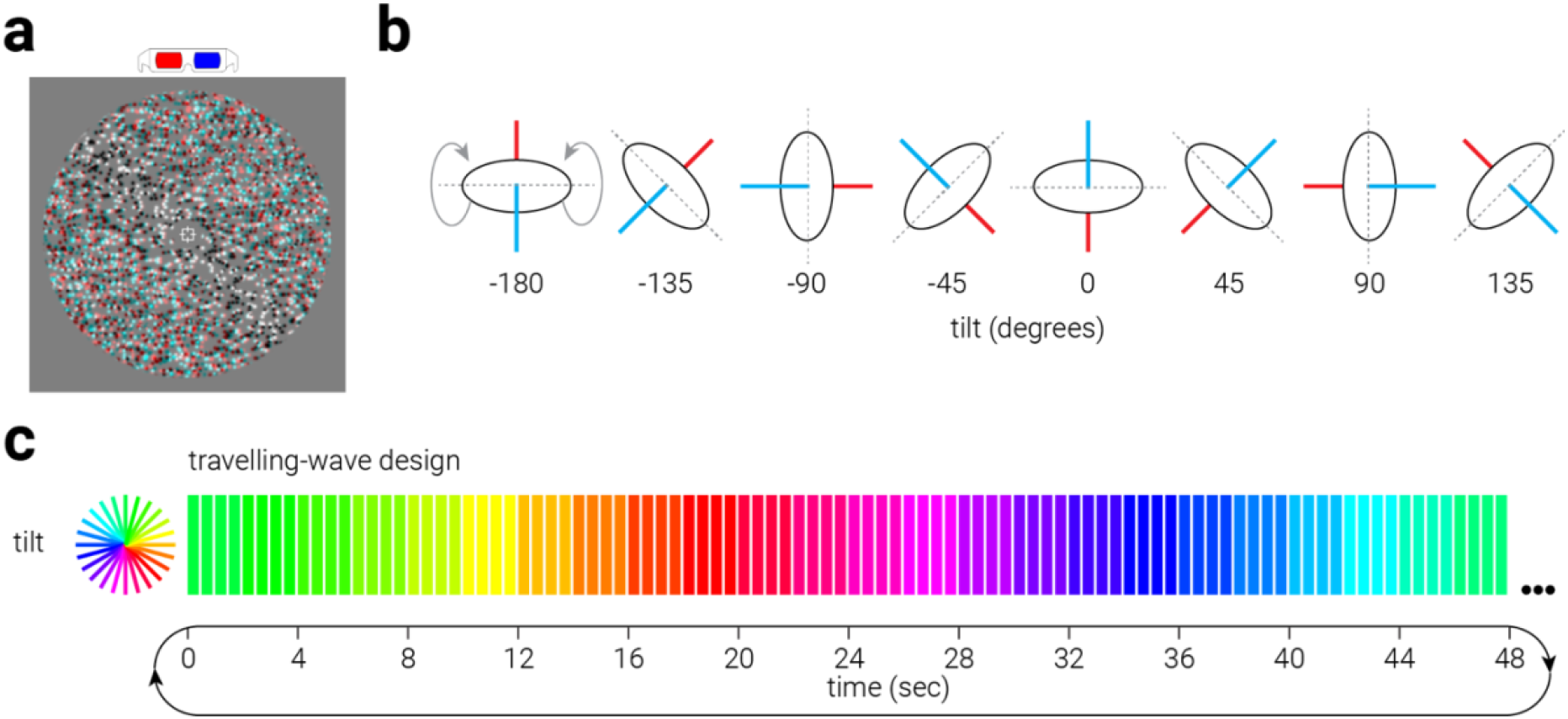
Three-dimensional stimuli and travelling wave procedure. **a**) Example random-dot stereogram stimuli used in the experiment (slant, −60°; tilt, 45°). **b**) Illustration of varying the tilt of a planar surface. **c**) Schematic diagram of travelling wave experimental design. A single cycle is shown, 7 cycles were repeated per run.

### Psychophysics procedure

We measured participants’ sensitivity to change in the tilt (at 60° slant) of the stimulus with a two-interval forced-choice experimental design in which the reference and test stimuli were presented in randomized order. The angular difference between test and reference stimulus was varied using the QUEST adaptive staircase method^23^. Each stimulus was presented for 500 ms with an inter-stimulus interval of 500 ms. Observers indicated which stimulus was tilted further clockwise using a keypress. We measured sensitivity to change around the four cardinal and four oblique angles. For each reference angle, we ran two randomly interleaved staircases, one that measured sensitivity to change in the positive direction and one that measured sensitivity to change in the negative direction; the final threshold estimate was derived from the average of the two staircases. Each staircase consisted of between 30 to 90 trials, depending on the number of trials required to reach a stable estimate. The order of reference angles tested for slant and tilt was randomized for each participant.

Observers could theoretically discriminate surface angle based only on the difference in depth at the edges (e.g., top and bottom) of a pair of stimuli. To minimize the availability of this cue, disparity-defined position was randomized by shifting the surface relative to the fixation plane (0° disparity) to between ±10% of the total surface depth.

### Travelling wave procedure

The slant angle of the plane was held constant 60° and we varied the tilt angle between 0-360° in 24 evenly spaced increments (**Fig. 1b,** right). A travelling-wave paradigm was designed to generate a traveling wave of activity across/within areas representing 3D orientation in visual cortex. This involved presenting planar surfaces which consecutively varied in tilt in either a clockwise or counterclockwise order. Planar surfaces at each tilt were presented for 2 s, with stimuli presented for 0.4 s separated by 0.1 s inter-stimulus-intervals consisting of only the background and fixation cross (gaps were used to reduce adaptation). Each cycle of stimulation lasted 48 s (**Fig. 1c**). fMRI runs consisted of 7 cycles in addition to a 16 s blank interval at the beginning and end of the run, thus taking 368 s. Each participant completed the experimental scanning session, which consisted of eight runs (four clockwise and four counterclockwise) and lasted approximately 60 min. Participant 1 (author RR) completed one additional session. This additional session was combined in the main analysis. We also collected data for variations of slant, but an artefact in the design of the slant travelling-wave procedure (but not the tilt procedure) meant that the data were uninterpretable and thus were omitted.

During the stimulus presentation, participants performed an attentionally demanding ring detection task. This served to ensured consistent attentional allocation on the stimulus during the experiment. Participants were instructed to fixate a central crosshair fixation marker and press a button when they detected a ring. Rings consisted of annular regions (width, 0.6°) of the stimulus, centred on fixation, that were reduced in contrast by 40%, presented at one of four possible eccentricities [1.8, 3.6, 5.4, 7.2]°. The shape of the ring presented to the left and right eyes was distorted to match the 3D orientation of the current planar surface. Rings were presented for the duration of the stimulus presentation (0.4 s) on 5% of presentations (selected pseudorandomly).

The method used to present images stereoscopically in the scanner prohibited eye tracking, so to assess eye movements in response to the stimuli we had participants complete four runs (two clockwise and two counterclockwise) of the travelling wave paradigm outside the scanner while we monitored their eye position.

### Magnetic resonance imaging

Data were collected at the Wolfson Brain Imaging Center with a 3T Siemens PRISMA system using a 32-channel head coil. Blood oxygen level-dependent (BOLD) functional data were acquired with a gradient echo-planar imaging sequence [echo time, 29 ms; repetition time, 2000 ms; voxel size, 1.5 × 1.5 × 2 mm; 30 slices covering occipital cortex] for experimental and localizer scans. Slice orientation was near to coronal section but rotated about the mediolateral axis with the ventral edge of the volume more anterior in order to ensure coverage of lateral occipital cortex (**Fig. 2a**). A T1-weighted, 3D-MPRAGE pulse sequence, anatomical scan (voxel size, 1 mm isotropic) was additionally acquired for each participant. These anatomical scans were used to register the functional data across scanning sessions, restrict the analysis of BOLD activity to grey matter voxels, and create flattened surfaces on which cortical activity is visualized.

**Figure 2.**
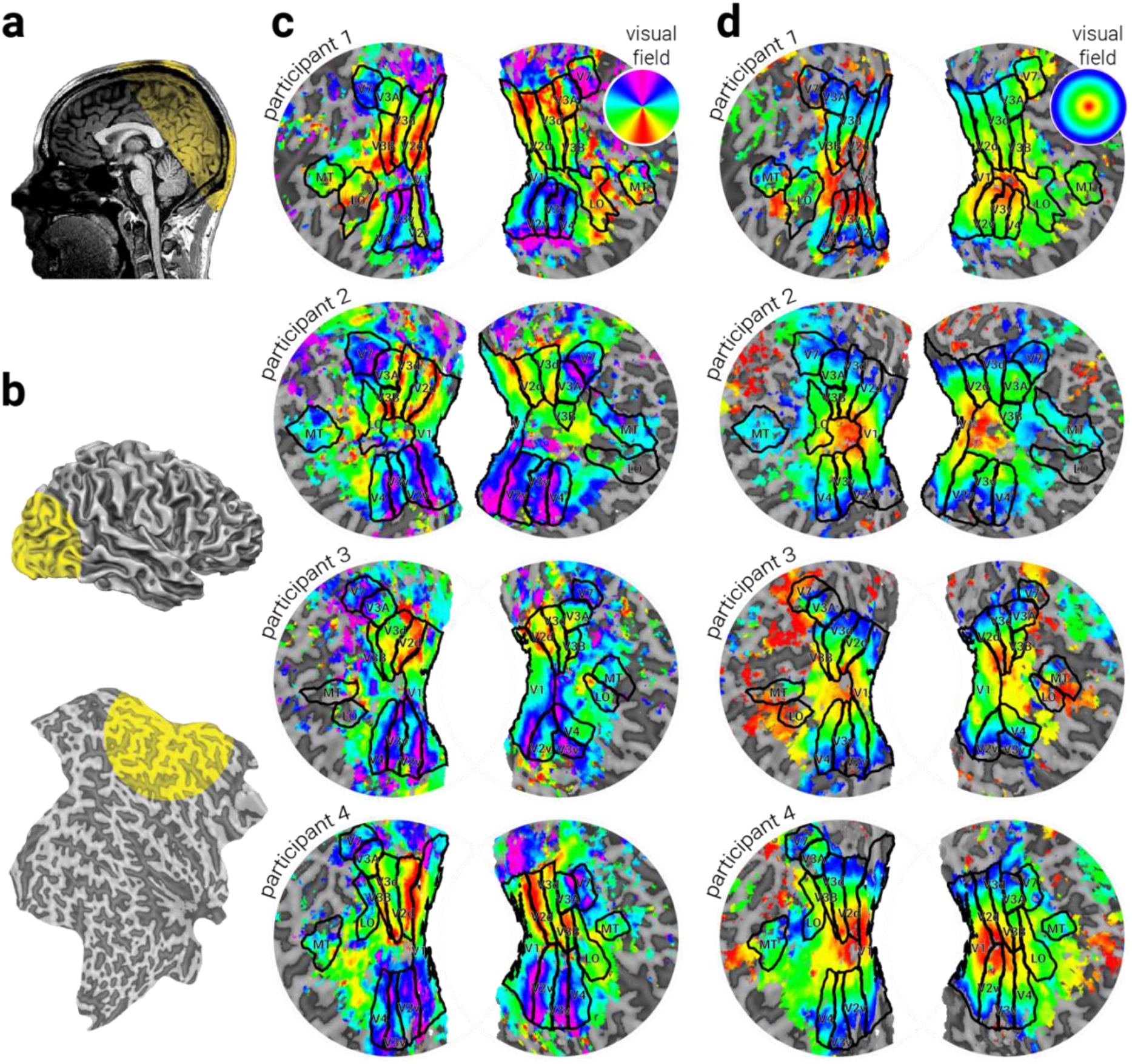
Retinotopic maps. **a**) Sagittal view of a structural MRI scan of participant 1, with coloured overlay illustrating slice orientation for functional scans. **b**) Reconstruction and flattened surface of structural scan, with coloured overlay illustrating circular regions shown in (**c-d**). (**c-d**) Flattened structural surfaces with coloured overlay illustrating phase map in response to **c**) a rotating wedge and **d**) expanding/contracting concentric rings. Black lines in (**c-d**) indicate retinotopic area demarcation.

### Data processing

Anatomical scans were aligned along the ACPC plane, grey matter and white matter were segmented, and flattened surfaces were computationally rendered with BrainVoyager QX (BrainInnovation, Maastricht, The Netherlands; **Fig. 2b**). Flattened surfaces (flat maps) were rendered such that areas that were concave on the original surface (i.e., sulci) are dark grey, and areas that were convex (i.e., gyri) are light grey. Functional data were preprocessed with slice timing correction, head motion correction, and high-pass filtering before being aligned to the participant’s anatomical scan. Functional images from the first run within a session were manually aligned to the anatomical, and these alignment coordinates were used to align the images from the remaining runs. Analysis of functional imaging data was performed in MATLAB (MathWorks, Natick, MA) using in-house scripts.

To evaluate periodic responses to 3D orientation, the data were analysed using a Fourier transformation-based method, well documented in the standard retinotopy studies^24–26^. As previously described by Ban et al. (2003)^27^, clockwise and counterclockwise run time series were concatenated and a discrete Fourier transformation was computed for each voxel after converting the raw signals to percentage signal changes. The phase measurement computed from this analysis was averaged between counterclockwise runs and reversed before being averaged with data from the clockwise runs to remove the effect of the hemodynamic delay in deriving the 3D orientation maps. Then, statistical testing to estimate the significance of correlation of BOLD signal at the stimulus frequency (1/48 Hz) was performed by comparing the squared amplitude at the stimulus frequency with the sum of squared amplitudes at the other frequencies, which yielded an *F*-ratio. The *F*-ratio was converted to a *P* value considering degrees of freedom of the signals (number of time points). In mapping retinotopic responses on the cortical surface, the polar angle phase of each significantly activated voxel was displayed using a continuous colour scale. We used a voxel-level Fourier *F*-statistic value (F(2, number of time points) > 7.38 and p < 0.05) as a criteria of significance for all Fourier-based analysis. Additionally, coherence values were obtained by calculating the voxel-by-voxel correlation between the concatenated time series and the corresponding best-fitting sinusoid at the frequency of the visual stimulation; coherence values were averaged between clockwise and counterclockwise runs. The coherence measures signal-to-noise^24,28^, and ranges from 0 to 1, where values near 1 indicate the fMRI signal modulation at the stimulus period is large relative to the noise (at the other frequency components) and values near 0 indicate that there is no signal modulation or that the signal is small compared with noise.

### Retinotopy

In a separate session, we performed ROI localization for each participant using retinotopic and stimulus contrast mapping procedures; the details of which are reproduced from Murphy, Ban, and Welchman (2013)^29^. Retinotopically organized visual areas (V1, V2, V3v, V4, V3D, V3A, V3B/KO, and V7) were defined using polar (**Fig. 2c**) and eccentricity maps (**Fig. 2d**), which were derived from fMRI responses to rotating wedge and expanding concentric ring stimulus presentations, respectively^25,26^. Area V3 was separated into ventral and dorsal quadrants in each hemisphere (V3v/d) consistent with previous delineation based on functional and cytoarchitectonic differences^30^. V4 was defined as the ventral region of visual cortex adjacent to V3v comprising a representation of the upper quadrant of the contralateral visual field^31,32^. V7 was defined as the region of retinotopic activity dorsal and anterior to V3A^3,31,32^. V3B/KO^4,33^ was defined as the area containing the union of retinotopically mapped V3B and the kinetic occipital region (KO), which was functionally localized by its preference for motion-defined boundaries compared with transparent motion of white and black dots^4,31,33,34^. This area contained a full hemifield representation, inferior to V7 and lateral to, and sharing a foveal confluence with, V3A^32^. Human middle temporal complex (hMT+/V5) was defined as the set of voxels in lateral temporal cortex that responded significantly more strongly to transparent dot motion compared with a static array of dots^35^. Lateral occipital complex (LOC) was defined as voxels in the lateral occipito-temporal cortex that preferred intact, compared with scrambled, images of objects^36^. LOC subregion LO was defined based on overlap of anatomical structures and functional activations, in line with previous work^37^.

### Pinwheel analysis

Previous work in which evidence for pinwheel-like encoding structures has been presented have relied on subjective identification of the formations. Here we developed a novel analysis method to objectively quantify the likelihood that the pattern of encoding at a particular location on the cortex is consistent with a pinwheel formation. To do so, we implemented a spotlight analysis of the flat map representations of tilt angle preference. The flat maps comprise polygons, each of which corresponds to a grey matter voxel from the anatomical image and has a phase value that indicates the tilt preference of that voxel. The phase values within the spotlight were compared to the position of each polygon centroid, relative to the center of the spotlight, in polar angle. A circular-circular regression between these values was calculated to determine their correspondence, independent of absolute phase. Regression values varied from 0 to 1, where 0 indicates no correspondence and 1 indicates perfect correspondence. Thus, if the phase values are structured in a pinwheel formation, their value will be predicted by the angular position from the center of the pinwheel, regardless of the phase offset position. The spotlight analysis was performed at each centroid location, with the coordinates of the centroid defining the center position of the spotlight. To reduce spurious correlations, only tests that included a minimum spotlight data point density of 1.4 (>216 data points) were included in the analysis. We present results obtained when the spotlight radius was 7 units in the flat map coordinate space (see the white circle in **Figure 7b** for size reference); however, to ensure that this did not bias our results, we systematically varied the spotlight radius between 2 and 14 units, finding a similar pattern of results.

## RESULTS

Following retinotopic mapping of visual areas (**Fig. 2c-d**) we measured participants’ phase-encoded responses to 3D tilt from 0-360° at 60° slant, defined by binocular disparity. **Figure 3a** shows the coherence of the activity measured from visual cortex with the frequency of the travelling wave stimulus. There was a relatively high coherence within retinoptopically defined visual areas, especially dorsal stream regions, suggesting that the neural populations within these regions were sensitive to aspects of the stimulus which varied as a function of tilt. **Figure 3b** shows the corresponding phase maps for each participant, which were thresholded to remove phase values with P>.05.

**Figure 3.**
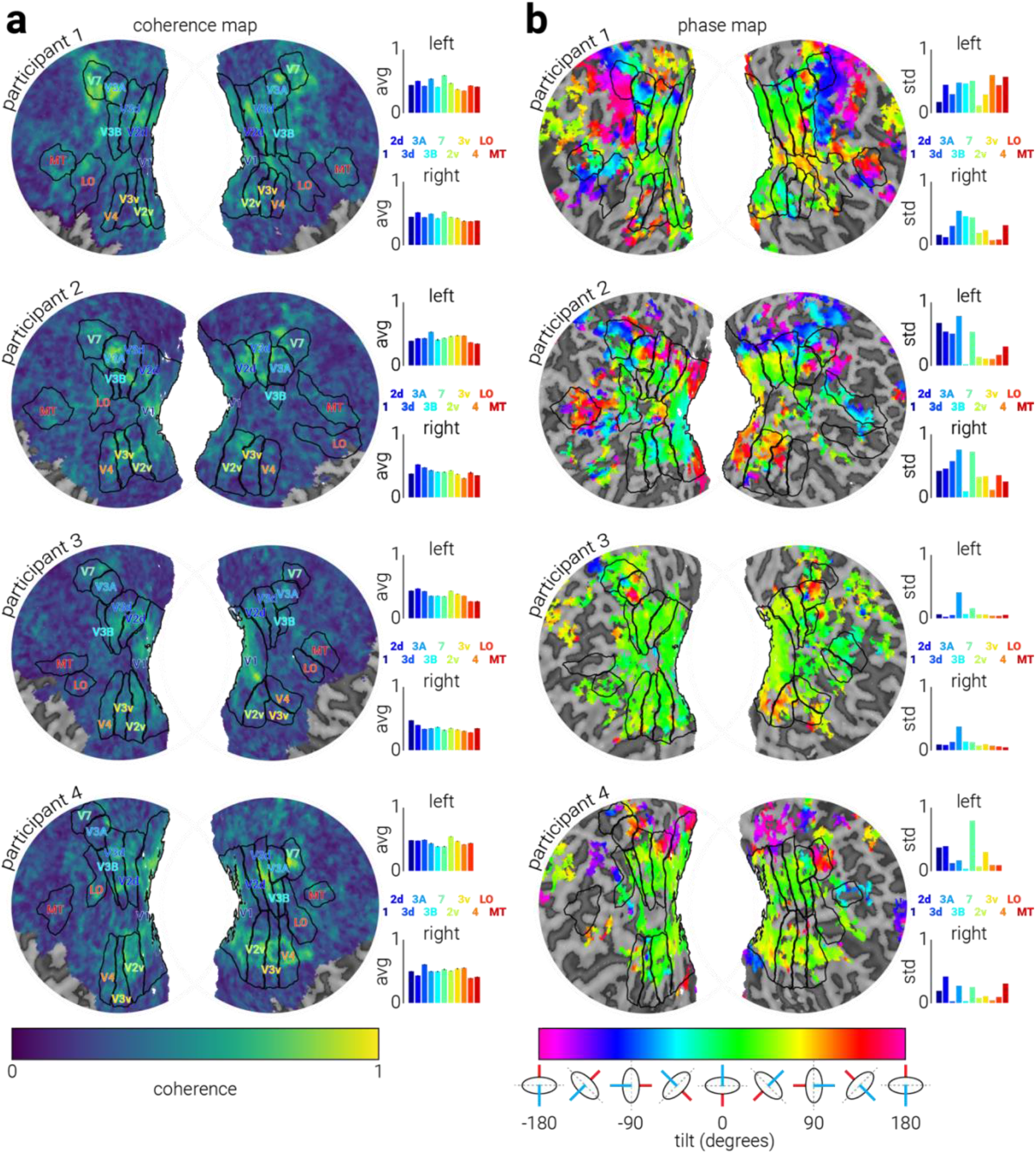
Cortical representation of tilt. Tilt **a**) coherence and **b**) phase maps for all subjects with retinotopic ROI borders overlaid in black. The plots on the right of the maps show the **a**) average coherence and **b**) standard deviation of phase values within each retinotopic area.

Summary analysis of voxel preference across the cortex revealed that participants 3 and 4 showed a high proportion of phase values corresponding to 0° tilt; however, this was only weakly present in participant 1 and not present in participant 2 (**Fig. 4a**). By contrast, all participants showed better performance for cardinal tilt angles than oblique angles (**Fig. 4b**). Previous work has identified areas V3A, V7, and MT as regions in which 3D orientation is encoded^10,12–14^. We reasoned that visual areas integral to encoding tilt would show selectivity to a variety of tilt angles. Thus, to assess the heterogeneity of tilt encoding, we calculated the variability of voxel phase preference within each visual area (**Fig. 4c**, right). In line with previous work, we found that areas V3A and V7 had more heterogeneous tilt preference than the other visual areas. By contrast, assessment of average coherence across visual areas suggested similar levels of coherence (**Fig. 4d**). A possible concern is that patterns of activity may have been influenced by eye movements during stimulus presentation. However, we found no evidence for this; in a separate session outside the scanner in which participants performed the same task, we found binocular eye movements were stable between different tilt angle presentations (**Supplementary Figure 1**).

**Figure 4.**
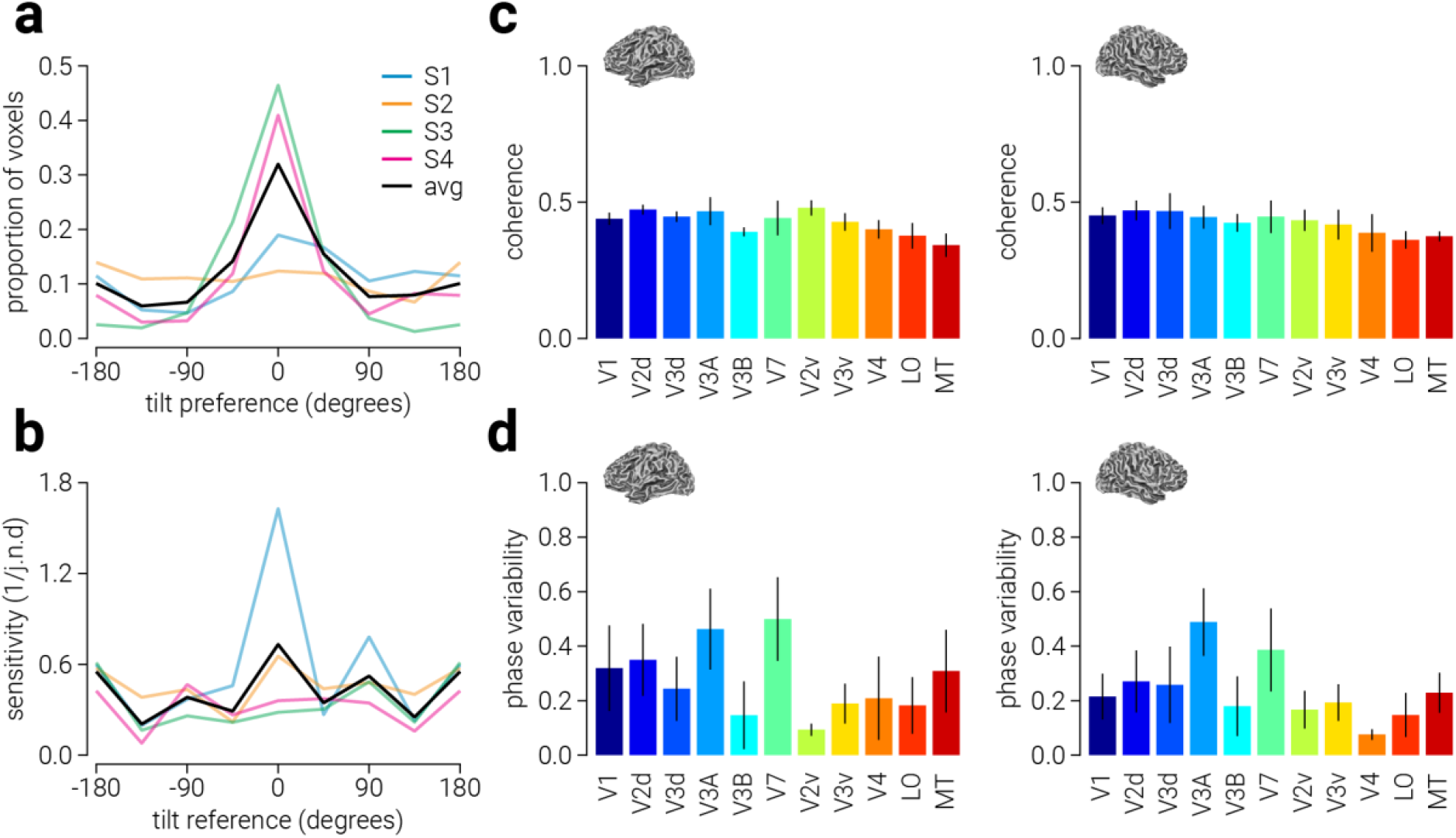
Summary of tilt representation. **a**) The proportion of significant voxels (P<.05) within the recorded region as a function of slant preference. **b**) Tilt sensitivity (reciprocal of the just noticeable difference) as a function of the reference slant angle. Colour labels in (**b**) match those in (**a**). (**c-d**) Average (**c**) coherence and (**d**) phase variability within retinotopic areas in the left and right hemisphere across all participants.

Another possibility is that the increased encoding variability observed in areas V3A and V7 is not specific to the representation of tilt in these areas. That is, information may be more heterogeneously encoded in these areas than other visual areas, regardless of the feature. To test this possibility, we performed the same analysis on the polar angle and eccentricity retinotopic data (**Figure 2b & c**); if all visual information is heterogeneously represented within V3A and V7, we would expect to find higher phase variability in these areas for polar angle and eccentricity. By contrast, the pattern of results for phase angle and eccentricity was markedly different than that for tilt angle. For polar angle, phase variability was highest in V1, and while it was second highest in V3A, it was relatively low in V7 (**Figure 5a**). For eccentricity, phase variability was also highest in V1, and relatively low in V3A and V7 (**Figure 5a**). These results indicate that the heterogeneous encoding of tilt angle in V3A and V7 was not simply due to these areas encoding all features in a varied manner.

**Figure 5.**
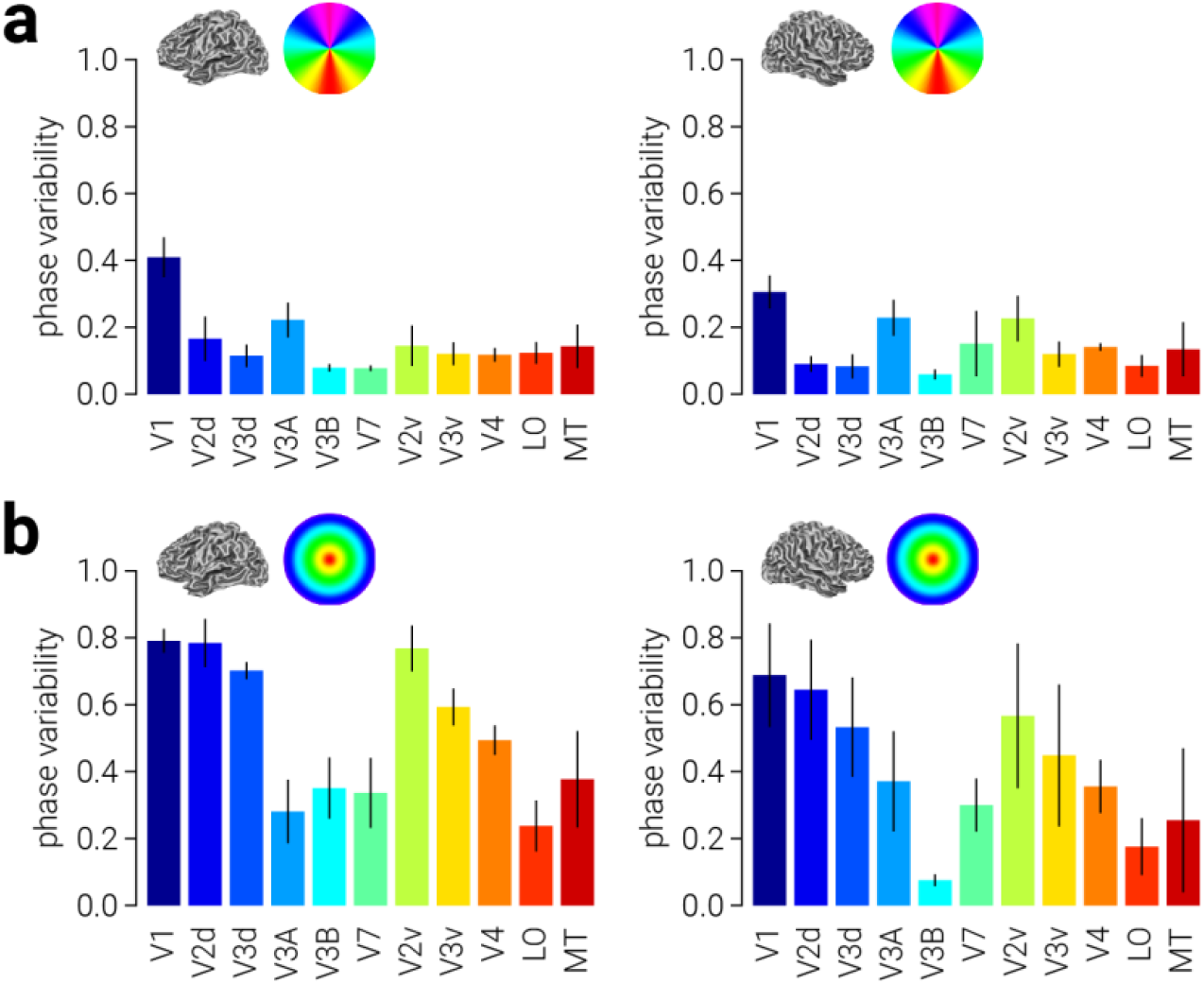
Summary of polar angle and eccentricity representation. Average phase variability for (**a**) slant angle and (**b**) eccentricity within retinotopic areas in the left and right hemisphere across all participants.

Inspection of areas V3A and V7 revealed two types of structure (**Fig. 6a**). Participant 2 had pinwheel-like structure of tilt preference in V3A (**Fig. 6b**, broken white circles). Participants 1 and 4 showed abrupt tilt preference reversal, between the border of V3A and V7 (**Fig. 6a**, broken white rectangles). With the exception of participant 1, these structures were not bilaterally present.

**Figure 6.**
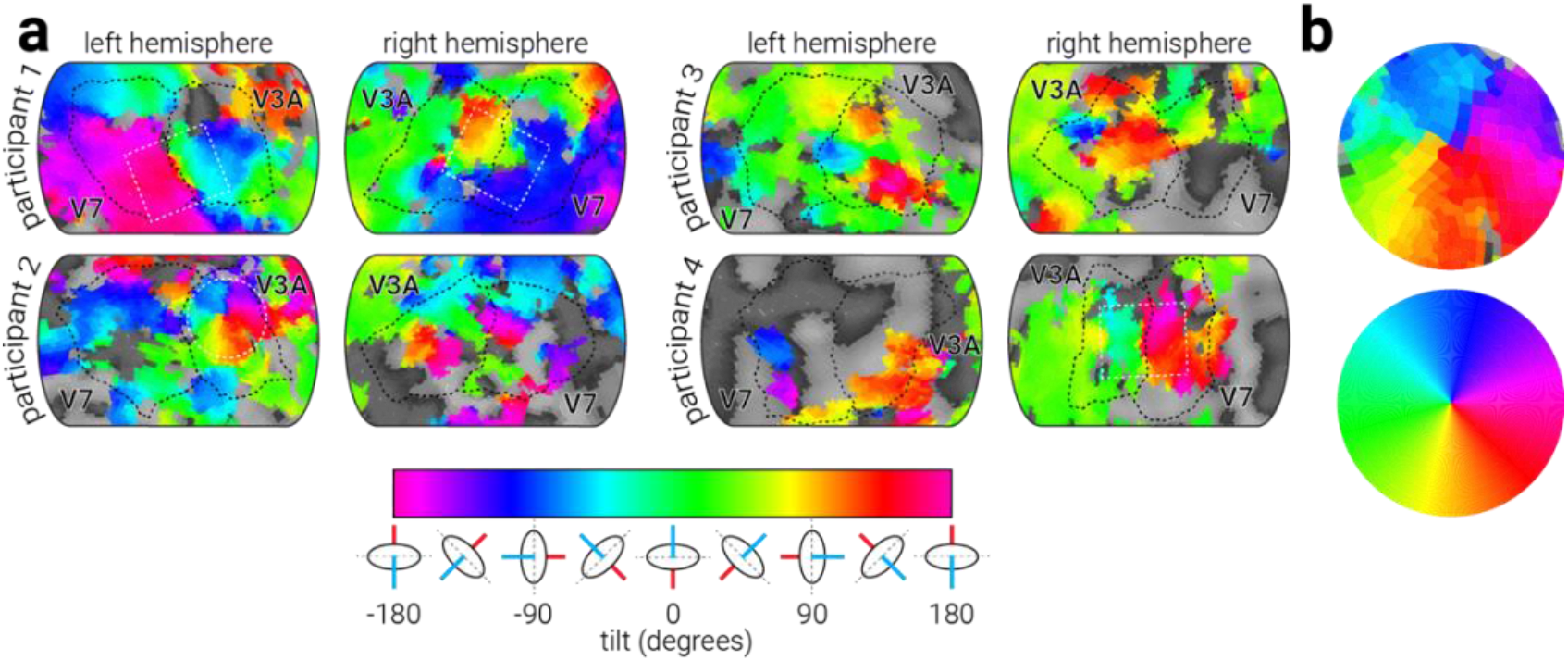
Representation of tilt in V3A/V7. **a**) Tilt phase maps of areas V3A and V7 for all participants. Broken black lines indicate region borders, broken white lines highlight pinwheel formation (circle) and intraregional phase transitions (rectangles). **b**) Comparison of the pinwheel like structure highlighted in the left hemisphere of participant 2 (top) and the tilt phase colour wheel (bottom).

It is possible that the pinwheel-like structure we observed in the left V3A of participant 2 was a product of chance. To test this possibility, we designed a pinwheel detection analysis that scans a map of phase values to yield a corresponding map of ρ values that indicate how well each location in the phase map corresponds to a pinwheel formation (see Methods for detailed explanation of the pinwheel analysis). As a sanity check, we first used the pinwheel detection analysis on an image of pinwheel-like structures in macaque V2 that correspond to motion direction preference^38^ (**Fig. 7a**, top). As expected, the corresponding correlation map shows peaks at locations corresponding to the centres of the pinwheel-like structures in the image (**Fig. 7a**, bottom). We then applied the analysis to participant 2’s right hemisphere phase map. We found that the location at which the highest correlation was found corresponded to the centre of the pinwheel structure in V3A (*ρ*=.917, (cos, sin) *P*=[1.1e_−5_, 3.6e_−6_]; **Fig. 7b**, top, white circle). A possible concern is that the pinwheel-like structure in V3A was a result of random distribution of phase values across the cortex, rather than a representing a functionally beneficial encoding structure. Given that the pinwheel is located in the visual area most intrinsically involved in processing 3D orientation (V3A), this seems unlikely. However, to test this possibility, we reran the pinwheel analysis on participant 2’s phase map after assigning phase values according to a two-dimensional pink noise (1/frequency) distribution (**Fig. 7b**, bottom). **Figure 7c** shows the distribution of correlation values from the pink noise simulation (pink bars) relative to the peak value found in the data (black arrow). The peak correlation value corresponding to the pinwheel structure in V3A clearly lies outside the values generated by pink noise, indicating that this kind of structure would be unlikely to be produced by chance.

**Figure 7.**
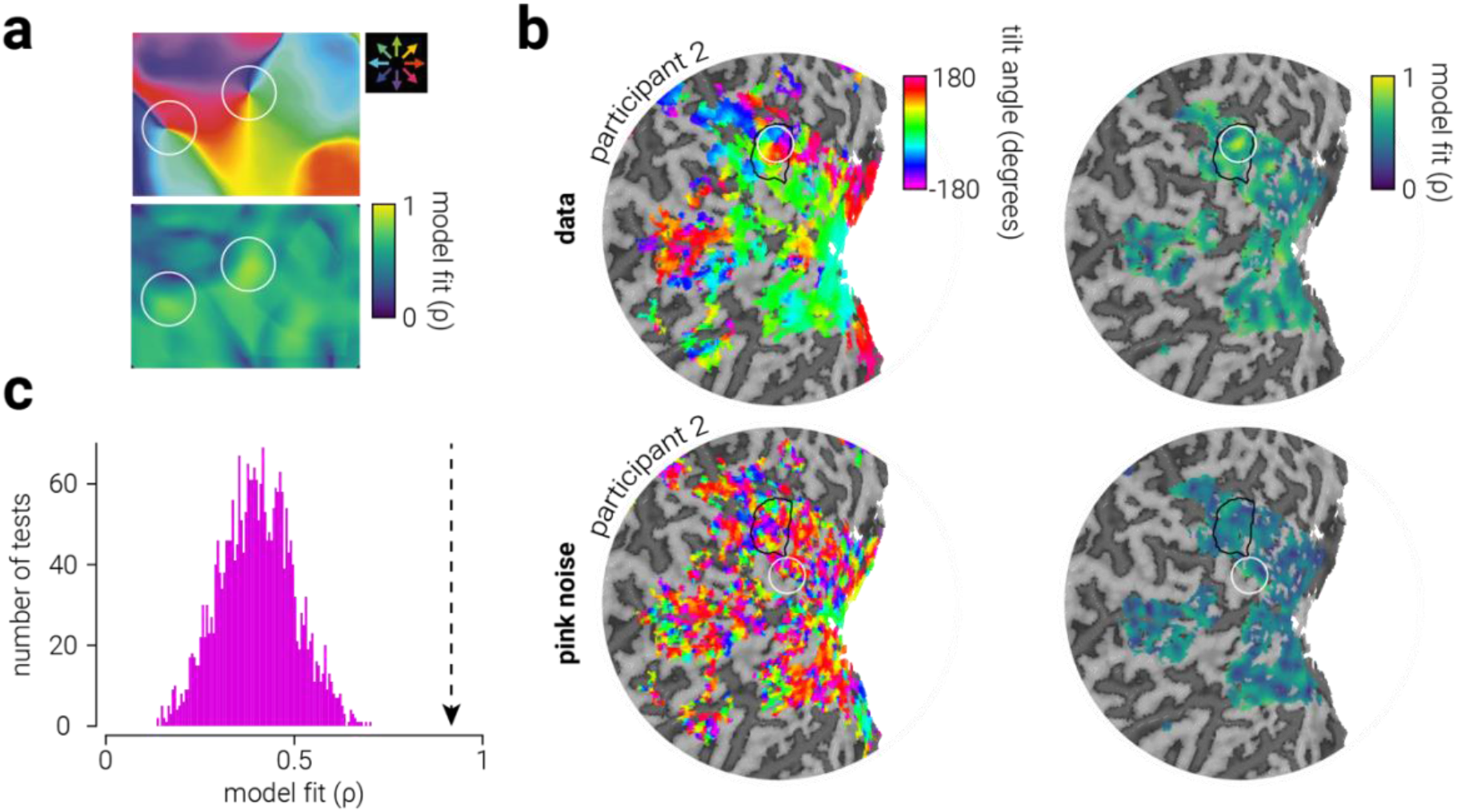
Pinwheel detection spotlight analysis. **a**, top) Motion direction preference map in macaque V2, shown using optical imaging^38^. **a**, bottom) Pinwheel correlation map produced by running the pinwheel detection spotlight analysis on (**a**, top). White circles indicate the size of the spotlight used in the analysis. **b**, top-left) Phase map and corresponding (**b**, top-right) pinwheel correlation map of Participant 2’s left hemisphere. The black outline delineates area V3A and the position and size of the white circle indicate the point of maximum correlation and the size of the spotlight, respectively. **b**, bottom) Same as (**b**, top), but when assigning phase according to a two-dimensional pink noise distribution. **c**) Histogram of the correlation values in (**b**, bottom-right); the peak correlation value in (**b**, top-right) is indicated by the arrow.

## DISCUSSION

Previous fMRI work with humans has identified areas V3A and V7 as responsive to 3D orientation^13,14^. These studies have employed experimental designs in which conditions are contrasted to reveal activity related to a particular cue. This procedure has been effective in isolating regions that support encoding of 3D orientation; however, it is blind to any putative functional organization of 3D orientation selectivity within these areas. Functional organization is likely to be the rule, not the exception, in the brain, as it has the potential to provide computational benefits that arbitrary organization cannot. There are many examples of functional organization within visual cortex at multiple scales. Classic examples include orientation/ocular dominance columns^39,40^ and retinotopic organization^25^. In a more recent example, relative depth was shown to be organized in area V3A_1_ in a way that persisted across scan sessions, although no identifiable functional structures were observed. Here we use phase-encoded presentation of planar surfaces with linear gradients of horizontal disparity to test whether the representation of 3D orientation is functionally organized within visual cortex. In particular, we measured fMRI activity of four participants to determine the preference of voxels within visual cortex for tilt angles between 0-360° with 60° slant angle.

We found a nonuniform distribution of tilt preference within visual cortex, there was a high proportion of voxels which preferred 0° tilt. Behaviourally, we found that observers were more sensitive to change in tilt angle around the cardinal axes, compared to the oblique. This is consistent with sensitivity to 2D orientation and motion, which is also best around cardinal axes (i.e., the oblique effect), and likely reflects an anisotropic distribution of 3D oriented surfaces in the environment. This may partially explain the anisotropic distribution of tilt preference observed. However, the large proportion of voxels that were identified as preferring 0° tilt may also reflect the anisotropic representation of near and far disparities in the upper and lower visual field. Perceptually, human observers are better at detecting targets defined by crossed (near) disparity^41^ and overestimate the distance of objects in the upper visual field^42–44^, while they are better at detecting targets defined by uncrossed (far) disparity and underestimate the distance of objects in the lower visual field^45^. This bias has also been observed physiologically: clustered regions that respond to near and far stimuli are preferentially located in retinotopic regions representing the upper and lower visual fields of V2/3, respectively^46^. Although all stimuli in the tilt condition had equal absolute global disparity, the anisotropic representation of near and far disparity in the upper and lower visual field likely resulted in a larger response to the 0° tilt stimuli, thus biasing the distribution of voxel tilt preferences.

We observed two patterns of functional organization of tilt preference in V3A/V7. The first, and perhaps most striking, was a pinwheel-like structure in the left hemisphere of participant 2. This structure encompassed the majority of V3A. Pinwheel-like functional organization has been observed for orientation selectivity in V1^19,20^. However, the spatial scale of these structures is very different; orientation pinwheels in V1 are ~300 μm in radius, whereas the tilt pinwheel we observed in V3A was ~10 mm in radius. In V1, this structure is thought to bestow at least two benefits for orientation processing: i) preferences for all orientations are brought together, within reach of upper-layer dendrites, and ii) orientation preferences on opposite sides are always perpendicular^20^. These same benefits may be shared by neurons that encode tilt in V3A; however, as tilt is a vector whose singularities (i.e., the point at the center of the pinwheel) rotate through 360°, tilt angles on opposite sides of the singularity are opposite, as opposed to perpendicular. While the benefits of pinwheel-like functional organization remain an empirical question, patterns resembling this formation that we observe here might suggest that pinwheels are used to represent a different visual property (3D tilt angle) in a different region (V3A) at a different spatial scale (~30 times larger). While our results are not fully conclusive in that these were not seen across hemispheres or in all participants, they might nevertheless hint that pinwheels represents a canonical organizational structure within the brain.

We found evidence of functional structures that were common to some, but not all participants. It is possible that these structures are not universal; however, it is also possible that we did not detect these structures across all participants due to insufficient resolution and/or signal strength. Future work could build on the findings revealed here by using fMRI at higher field strength^47^ (e.g., 7T) to target and characterize the representation of 3D surface orientation in areas V3A and V7. Further, although we used binocular disparity to define the planar surfaces in the current study, V3A and V7 have been shown to respond to 3D orientation defined by motion^11,15,16^, texture^17^, and shading^18^. Thus, future work could compare the representation of 3D orientation within these areas using different depth cues in isolation and combination.

Using phase-encoded presentation of 3D stimuli we find evidence in support of the functional organization of 3D orientation within visual cortex. In one participant we find fMRI response patterns suggesting that tilt preference is unilaterally organized in a pinwheel-like structure, similar to those observed for orientation preference in V1^19,20^, that encompasses most of area V3A. Further, for two participants we find an abrupt reversal in tilt preference at the (retinotopically defined) border between V3A and V7. These findings indicate that 3D orientation is functionally organized within the human visual cortex, and the discovery of a large pinwheel-like structure suggests that this type of organization may be applied canonically within the brain at multiple scales.

## Acknowledgements

We would like to thank Hiroshi Ban for their help and insight on the project. The work was supported by the Leverhulme Trust (ECF-2017-573), the Issac Newton Trust (17.08(o)), and the Wellcome Trust (095183/Z/10/Z).

