## Supplementary Figure 1 for "Testing for functional organization of three-dimensional surface tilt encoding within visual cortex"

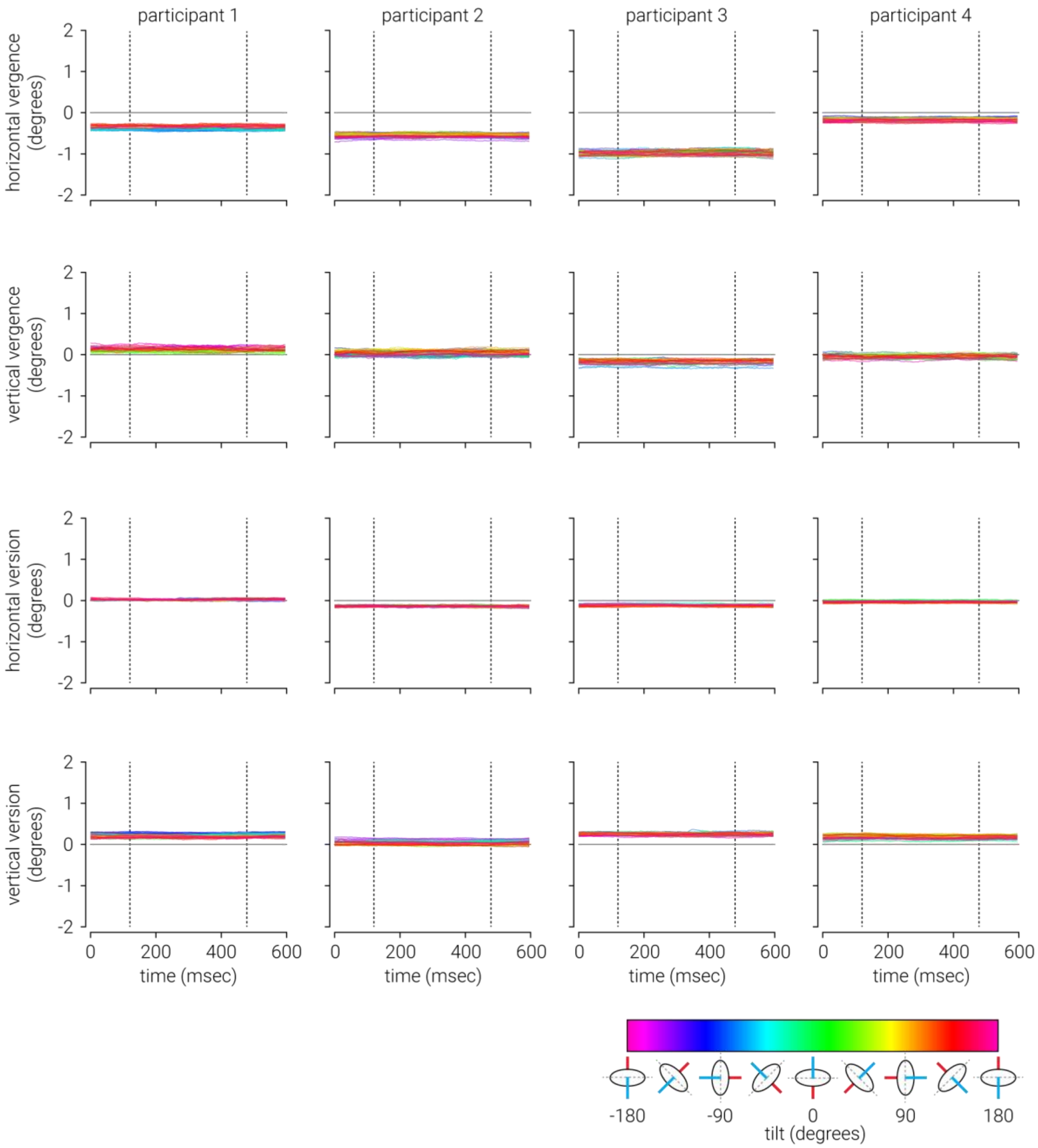


**Supplementary Figure 1. Eye position during presentation of tilt stimuli.** Average binocular fixation position in terms of horizontal/vertical vergence and version as a function of tilt for each participant during control runs of the travelling wave procedure conducted outside the scanner. The colour of each position trace indicates the corresponding tilt of the stimuli. The vertical broken black lines indicate stimulus on and off times.
